# Analysis of rhizosphere fungal community of agricultural crops cultivated in laboratory experiments on Chernevaya taiga soil

**DOI:** 10.1101/2023.09.14.557691

**Authors:** Irina Kravchenko, Mikhail Rayko, Sophie Sokornova, Ekaterina Tikhonova, Aleksey Konopkin, Alla Lapidus

## Abstract

Chernevaya taiga in Western Siberia, Russia, is a special environment with fertile soil, extraordinarily high herbaceous plant sizes, and incredibly quick rates of plant residues breakdown. We postulated that a unique rhizospheric fungal community would be created when growing crops on chernevaya soil, which has never been used for agriculture. These may be the source for the novel effective biostimulator and biocontrol fungal agents for modern agriculture. In this study, we analyzed the fungal communities in the rhizosphere of spring wheat and radish cultivated in greenhouse experiments on chernevaya and control soils using high-throughput ITS sequencing. Additionally reprehensive fungal strains were isolated and examined for the stimulation of wheat seedlings growth. The research shows that Ascomycota and Mortierellomycota were the most abundant phyla in the rhizospheric fungal community, mainly composed of *Mortierella* species. The control soils were abundant in Mucoromycota, represented mostly by *Mucor* and *Umbelopsis*. Potentially plant-pathogenic fungi *Fusarium and Oidiodendron* were found only in the rhizosphere of crops grown in the control soil. Whereas plants grown on chernevaya soil have a wide variety of possible biocontrol fungi. Novel fungal isolates demonstrated stimulating effect on the growth and biomass accumulation of wheat seedlings. The findings of the study imply that the creation of biorational products for crop growth and protection may depend on the taxonomic composition analysis of the rhizospheric mycobiota of the crops growing on the soil with the unique characteristics.

## Introduction

Chernevaya taiga soils offer an interesting and understudied subject for metagenomic research due to their tall grasses, low humus concentration, and very high plant residue decomposition rates. Furthermore, these soils could yield plant growth promoting microorganisms, such as cellulolytic strains and biologically active compounds, with practical applications. This presents an opportunity for exploring the bacterial and fungal communities of these soils, and their impact on soil productivity. Recently, we have investigated the rhizosphere bacterial microbiota and found that agricultural crops grown in laboratory conditions on chernevaya soil demonstrate specific rhizosphere microbial community’s composition (Kravchenko et al. 2022).

The root-associated microbiota influenced by biotic and abiotic factors in the soil and host plant (Munir et al. 2022). The rhizosphere (the area of soil in contact with plant roots) is rich in microbial diversity and interactions. Root exudates can influence composition and functions of microbial communities in the rhizosphere (Berg and Smalla 2009). Fungal interactions in the rhizosphere can be mutually beneficial, promoting nutrient acquisition and stress tolerance in plants (Philippot et al. 2013). Mycorrhizal, biocontrol, or mycoparasitic fungi have been studied for their positive effects on plant growth and health (Adedayo and Babalola 2023). Plant-growth-promoting fungi (PGPF) belong to various genera, such as *Mucor, Trichoderma, Cladosporium, Sordaria, Penicillium* etc. PGPFs can improve plant health by phytohormone production, phosphate solubilization, mitigation of stress, volatile organic compounds, etc. Under certain conditions, they are able to change physiological features of bacteria in the soil (Akinola et al. 2020). For example, the symbiosis of arbuscular mycorrhizal fungi (AMF) with mycorhizal helper bacteria (MHB) are comparatively well studied (Zhang et al. 2021).

Root-associated fungi, which make up a significant percentage of the root microbial community and can be symbiotic or pathogenic, can interact with plants (Baron and Rigobelo 2022). It is therefore of significant interest to comprehend how interactions between root-associated fungal communities may be related to plant health and agricultural productivity. Although many studies have focused on the bacterial composition of root-associated microbiomes (Faist et al. 2023; Nanetti et al. 2023; Zhang et al. 2020), while few have focused on fungi. Plant pathogens interact closely with the other members of the microbial community in the rhizosphere, and this interaction is crucial for the likelihood of plant infection. In order to manage the health and production of the plant, it is crucial to comprehend the variables that affect the composition and intermicrobial interactions of the rhizosphere mycobiome.

In a recent study, the mycobiome of bulk soil, rhizosphere soil, and root samples of soybean plants were examined. The results showed that the communities in high and poor yield sites had different compositions. The roots of poor yield locations included a higher relative number of ASVs associated with the well-known soybean diseases *Fusarium* and *Macrophomina phaseolina*. In contrast, the high yield sites exhibited a high abundance of the well-known plant-beneficial genera *Trichoderma* and *Metarhizium* (Bandara et al. 2021). It was shown that the makeup of various fungal communities in the rhizospheres of agricultural crops was altered by the use of mineral and organic fertilizers. According to Semenov et al.2022, fresh cattle manure used as organic fertilizer significantly enhanced fungal abundance while reducing fungal variety. Mineral fertilization (NPK) considerably boosted the relative abundances of fungal phytopathogens like *Alternaria* and *Fusarium*.

We hypothesize that the composition of rhizosphere fungal communities will have a strong correlation with crop productivity and that there will be crop-specific differences in community composition and indicator taxa for Chernevaya taiga sites.

The objectives of our research were; (i) to study the rhizosphere mycobiome of radish and wheat plants grown in plant cultivation set up in a laboratory on the chernevaya and control soil; (ii) to investigate differences in root mycobiome between samples from high and low productivity sites; (iii) to assess crop-specific differences in fungal community composition; and (iv) to obtain isolates of dominant rhizospheric fungi and test their effect on the growth of wheat seedlings.

## Materials and methods

### Soil characteristics and pot experiments

Pot experiments were conducted with the chernevaya soil—dark gray soil (Umbrisol, Albic, Loamic, Folic WRB classification) collected under the tallgrass fir-aspen forest with a shrinking fir stand (56.30693 N, 85.47063 E, hereinafter - T1), and the control soil - oligotrophic sandy loam soil (Retisol, Luvic, Folic WRB Classification) collected under the pine forest with an admixture of larch (56.48106 N, 84.79860 E, hereinafter - T3). The T3 site was chosen as a control, because it is located in close proximity and has a similar composition of vegetation. However, unlike the T1 site, there are no signs of plant gigantism and extremely high plant productivity. The soil sampling in the field was conducted in May 2020 on the territory of the Tomsk region, Russia. In previous publications, we have described in detail the areas and sites of study (Abakumov et al. 2020; Polyakov et al. 2022).

The pot experiment was conducted on 15 February-15 March 2021 as described in our previous publication (Kravchenko et al. 2022). Briefly, surface-sterilized seeds of radish (*Raphanus raphanistrum* subsp. Sativus, cultivar “Red light”) and spring wheat (*Triticum aestivum* L., cultivar “Lada’’) were germinated on moistened filter paper for 2 days. Then ten seedlings were transferred to pot contained 0.2 kg either T1 or T3 non-sterile soil with three replicates for each soil type. The experiment was carried out in a phytotron at 23 ± 1°C, 50–60% RH, 550 M photon/m^2^× s, and a 16 h photoperiod. The soil was kept at 60% water holding capacity by daily weighing and watering.

Plants were destructively sampled at the end of the experiment for rhizosphere microbial investigation. To extract DNA and isolate fungus, a soil fraction that was closely connected to the root system was taken.

### DNA isolation, amplification, and Illumina ITS rRNA sequencing

Total soil DNA was extracted from 0.25 g of rhizosphere soil samples using the Power Soil DNA Isolation Kit (Qiagen, Calsbad, CA, USA). The NanoDrop 1000 spectrophotometer (Thermo Scientific, Waltham, MA, USA) was used to determine DNA concentration and purity.

Fungal DNA was amplified according to the EMP ITS Illumina amplification protocol (Smith et al, 2016). Finally, the prepared DNA libraries were sequenced using an Illumina MySeq (Illumina, USA) in paired-end mode using the equipment of the resource center “Genomic Technologies, Proteomics, and Cell Biology” of ARRIAM.

### Bioinformatics analysis

The analysis was performed as described in our previous publications (Rayko et al 2021; Kravchenko et al. 2022). The Cutadapt software (Martin 2011), the Dada2 package (Callahan et al. 2016) included in the QIIME2 package, v-2022.8 (Bolyen et al. 2019) were used in the preprocessing of the raw Illumina reads (denoising, merging paired reads, and eliminating chimeras).

After filtering, high-quality amplicon sequence variants (ASVs) were obtained and used for subsequent analyses. The taxonomic classification of the obtained ASVs was performed using UNITE database v. 8.99 (Nilsson et al. 2019) as a pre-trained classifier for QIIME2. Then all ASVs not belonging to the Fungi kingdom were removed.

The ASV abundance values in each sample were standardized by using the lowest level of sequence coverage as a reference. To remove possible PCR artifacts or chimeras, only ASVs with an abundance of more than 4 reads and presented in at least 10% of samples were selected for subsequent analysis.

The phylogenetic tree of the obtained ASVs was inferred using the FastTree algorithm (Price et al. 2010). The package Microbiome Analyst (Dhariwal et al. 2017) was used to calculate alpha-diversity (Chao1 index) and beta-diversity (UniFrac, Bray–Curtis dissimilarity) metrics, and dimensionality reduction (PCoA, Ward’s hierarchical clustering) (Ward 1963).

LEfSe (Linear discriminant analysis Effect Size) approach was applied to identify differently abundant taxa (Segata et al. 2011). The threshold for the log2 fold change (l2FC) was set at 2.0, and the FDR-adjusted p-value was cutoff at 0.05 steps. The association of the fungal taxa with ecological guilds was performed using the FUNGuild approach (Nguyen et al. 2016).

### Isolation and characterization of rhizospheric fungi

By creating a decimal dilution from the soil samples from the rhizosphere, fungal strains were isolated. In order to stop bacterial growth, 0.05 mL of each dilution was added to Petri plates containing potato dextrose agar (PDA) and chloramphenicol. The dishes were then incubated at 25°C for 5–10 days. First, the morphological characteristics of the fungal isolates were looked at, including color, colony surface, colony margin, texture, and pigment secreted. To produce axenic cultures, colonies with various morphotypes were chosen and streaked repeatedly. First, fungal isolates were identified using morphological and cultural characteristics, including colony development pattern, conidial morphology, and pigmentation, using the identification keys (Domsch et al. 2007). The current names and positions of fungi were corrected using the Index Fungorum database (http://speciesfungorum.org/Names/fundic.asp).

One fungal isolate from each morphotype was sub-cultured for DNA extraction. The total genomic DNA was extracted from fresh fungal mycelia using a method (Boulygina et al. 2002) with the addition of 20-50 units of zymolyase enzyme per sample. Extracted DNA was amplified by PCR of the ribosomal operon ITS1 – 5.8 S – ITS2 - rRNA with the ITS4-ITS5 primer system (Angelov et al. 2015). The amplified PCR products were sequenced by the Sanger method at the Core Facility “Bioengineering” in the Research Center of Biotechnology RAS.

The obtained nucleotide sequences were aligned and compared to sequences available at the NCBI using BLASTN searches. Results from BLAST were compared with morphological identification. Specimens of each isolate are available in Core Facility “UNIQEM collection” at the Research Center of Biotechnology RAS, Moscow, Russia.

The five chosen fungal isolates were used to assess how each one affected the growth of the spring wheat seedlings. The seeds were prepared and allowed to germinate as described above. From 10-day-old cultures grown on PDA medium, 10^7^ spores/mL fungal conidia suspensions were prepared. Ten milliliters of the conidia suspension from each test isolate were mixed into the soil around wheat seedlings before they were transplanted into pots filled with unsterilized agricultural soil from the Moscow area. As a control, sterile water inoculation was used. For each treatment, 30 plants (n = 30) were employed, and three separate repetitions were carried out.

After 30 days, the seedlings were removed from the pots. The PGP ability of the fungal isolates was determined by measuring the seedlings’ height, root length, and dry and fresh weights. Dry weights were measured by drying in an oven at 65 °C for 24 h. The growth promotion effectiveness of fungi (FGPE) was assessed using the following formula:

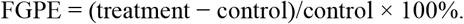

### Nucleotide sequence accession numbers

All the raw reads, commands and parameters used in data analysis are openly available in FigShare at https://doi.org/10.6084/m9.figshare.22639870.v1 (accessed on 16 April 2023).

Nucleotide sequences of the fungal isolates have been deposited in GenBank (OQ621732.1-OQ621735.1

## Results

### Fungal community structure and composition

In total, samples that were analyzed revealed representatives of seven phyla. Members of Kickxellomycota and Chytridiomycota were only found in control soils, whereas Rozellomycota and Chytridiomycota were only found in chernevaya taiga soils (Table 1).

**Table 1.** Actual phylum-level abundance of fungal amplified sequence variants (ASVs) connected with the rhizosphere of growing wheat and radish on control and chernevaya soils.

| Phylum | Control T3 soil | Chernevaya T1 soil |
| --- | --- | --- |
| Ascomycota | 11047 | 17781 |
| Basidiomycota | 4576 | 4080 |
| Chytridiomycota | 0 | 483 |
| Kickxellomycota | 26 | 0 |
| Mortierellomycota | 68 | 8010 |
| Mucoromycota | 31930 | 2786 |
| Rozellomycota | 0 | 265 |

Taxonomic analysis of the samples demonstrated that Ascomycota and Mortierellomycota were the most abundant phyla in the chernevaya T1 soil samples (Fig. 1). We earlier observed a high abundance of Mortierellomycota, represented mostly by the *Mortierella* genus, in the bulk soil samples from Chernevaya taiga (Rayko et al. 2021).The control T3 soils were abundant in Mucoromycota, represented mostly by *Mucor* and *Umbelopsis*.

**Fig. 1.**
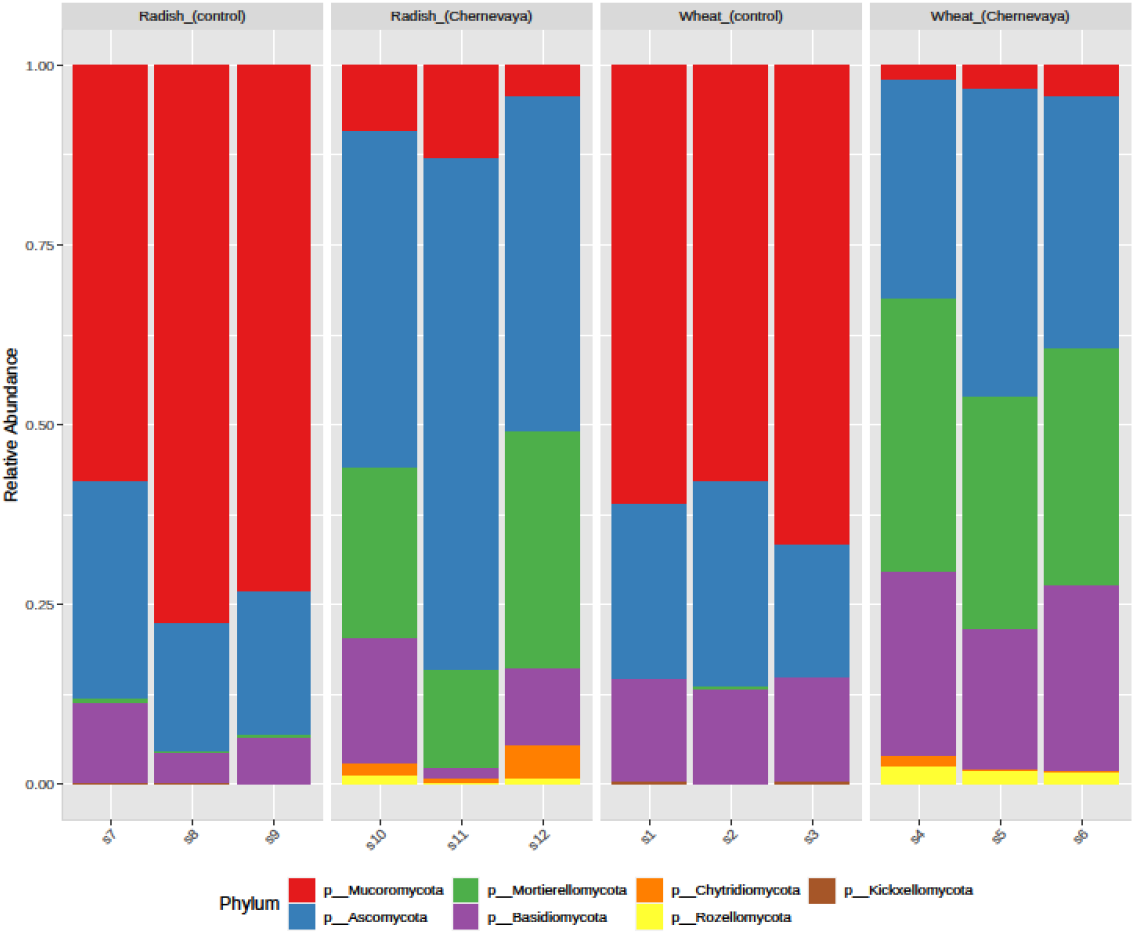
Actual abundance of fungal amplified sequence variants (ASVs) associated with the rhizosphere of radish and wheat plants grown on control and chernevaya soils at the phylum level

The differences in fungal rhizosphere communities were more significant among plants growing in chernevaya and control soils than among different plants (radish and wheat) growing on the same soil.

### Indicator Taxa

We clustered samples based on taxonomic composition and combined them on order level (sum up all the members of the corresponding order among ASVs) using the Ward method (Fig. 2). Grouping the samples according to soil type revealed a set of orders that shared these clusters.

**Fig. 2.**
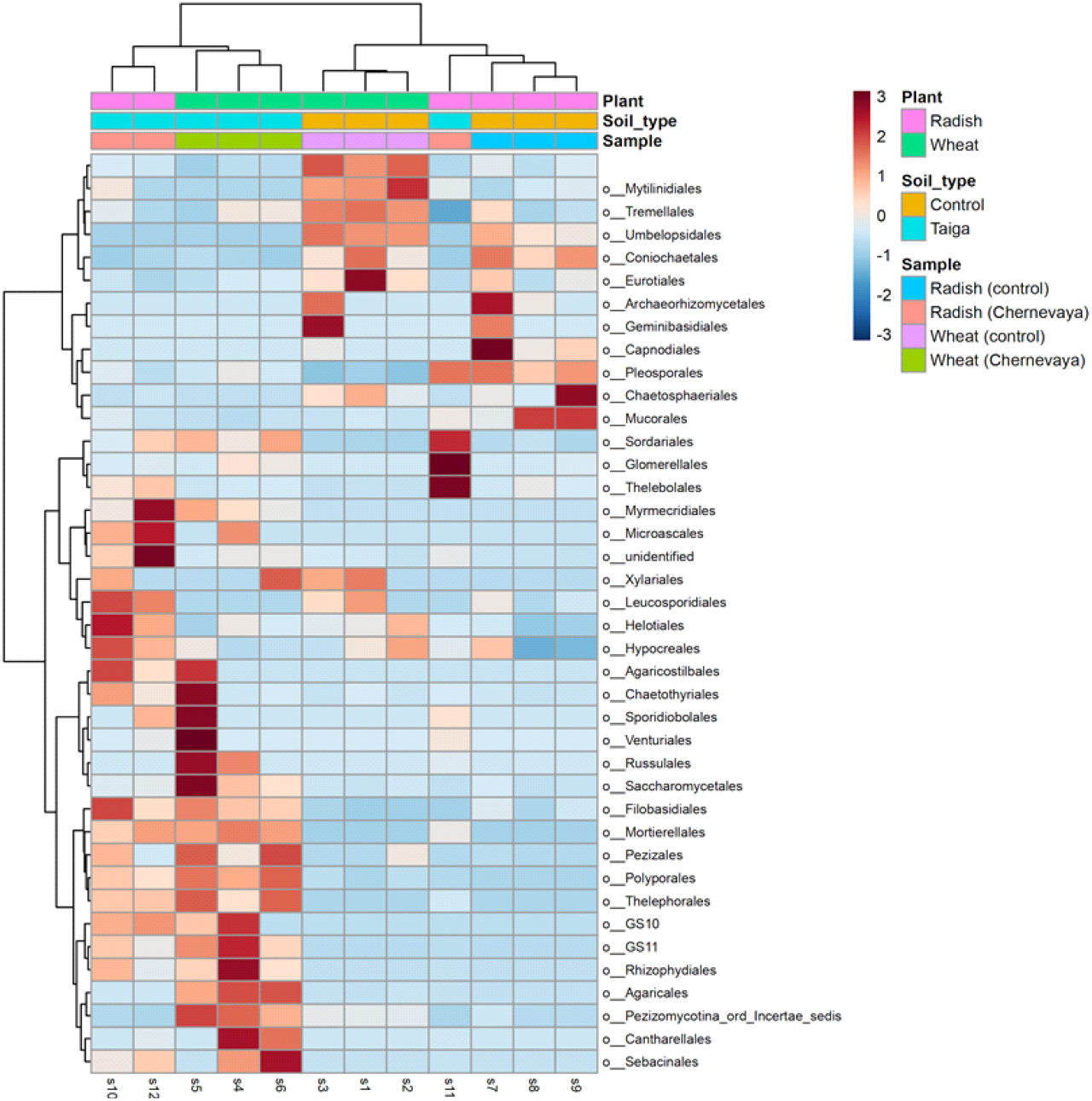
Heatmap visualization of clustering of fungal communities from the rhizosphere of the radish and wheat plants, grown in the chernevaya and control soils, based on the order relative abundance (Ward’s method). Columns represent individual samples, gradient - normalised relative abundance

Heatmap visualization reveals differences in the composition of fungal community both between different types of soils (T1, Chernevaya taiga soil, and T3, control) and types of plants. For example, in wheat rhizosphere we detected prevalence of the ascomycetous fungi Chaetothyriales and Glomerellales. Chaetothyriales are phenotypically plastic, black yeast-like fungi that are known as opportunistic pathogens of vertebrates and are frequently found as saprophytes. Glomerellales demonstrate very flexible ecological traits and include saprobes, endophytes, and phytopathogens. Glomerellales demonstrate very flexible ecological traits, and include saprobes, endophytes and phytopathogens.

Also, in the wheat rhizosphere samples from Chernevaya soils, one can see the presence of the Sebacinales. These fungi are related to Agaricomycetes, have a widespread distribution, and are mostly terrestrial. Many of them form mycorrhizae with a wide variety of plants.

We applied linear discriminant analysis to determine the log10 LDA score for each taxon in each sample in order to assess fungal taxa that might be less prevalent but have a significant impact on plant growth (Fig. 3). The most strongly selected or depleted taxa are usually mostly responsible for the differences in the rhizosphere community structure between the chernevaya and control soils.

**Fig. 3.**
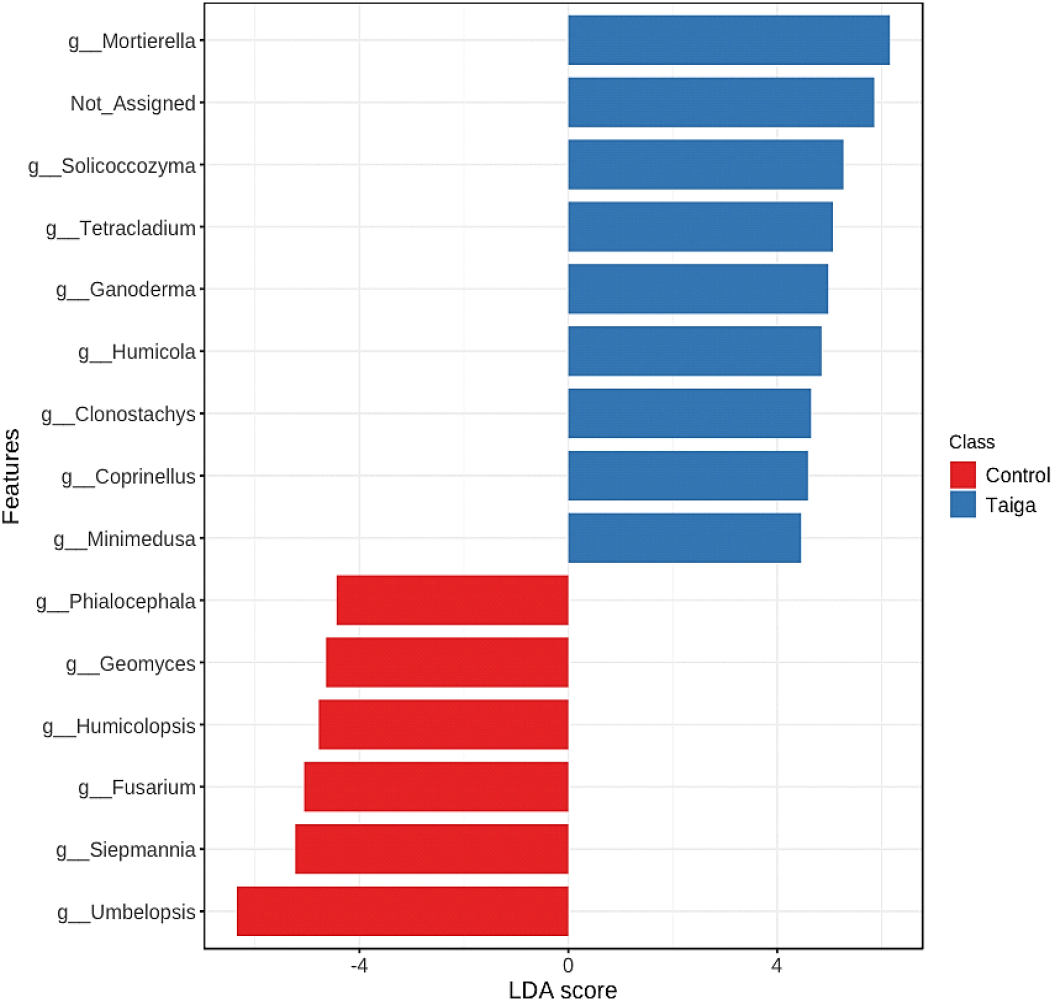
Effect size (LDA) for the fungal species in the rhizosphere of plants growing in the chernevaya soil relative to those in the control soil

LDA analysis showed that the rhizosphere of crops in Chernevaya taiga soils is characterized by representatives of various divisions: *Mortierella, Solicoccozyma, Tetracladiu*m. For control soil, another set of chracteristics was identified, which included *Phialocephala, Geomyces, Humicolopsis*, and *Fusarium*.

### Diversity analysis

We investigated the impact of soil type and host plant on the diversity of the fungal communities on Chao1 diversity index. Our results indicate a significantly higher alpha-diversity of the rhizospheric communities in the Chernevaya taiga samples both, assessed by t-test, p-value: 0.0095 (Fig. 4).

**Fig. 4.**
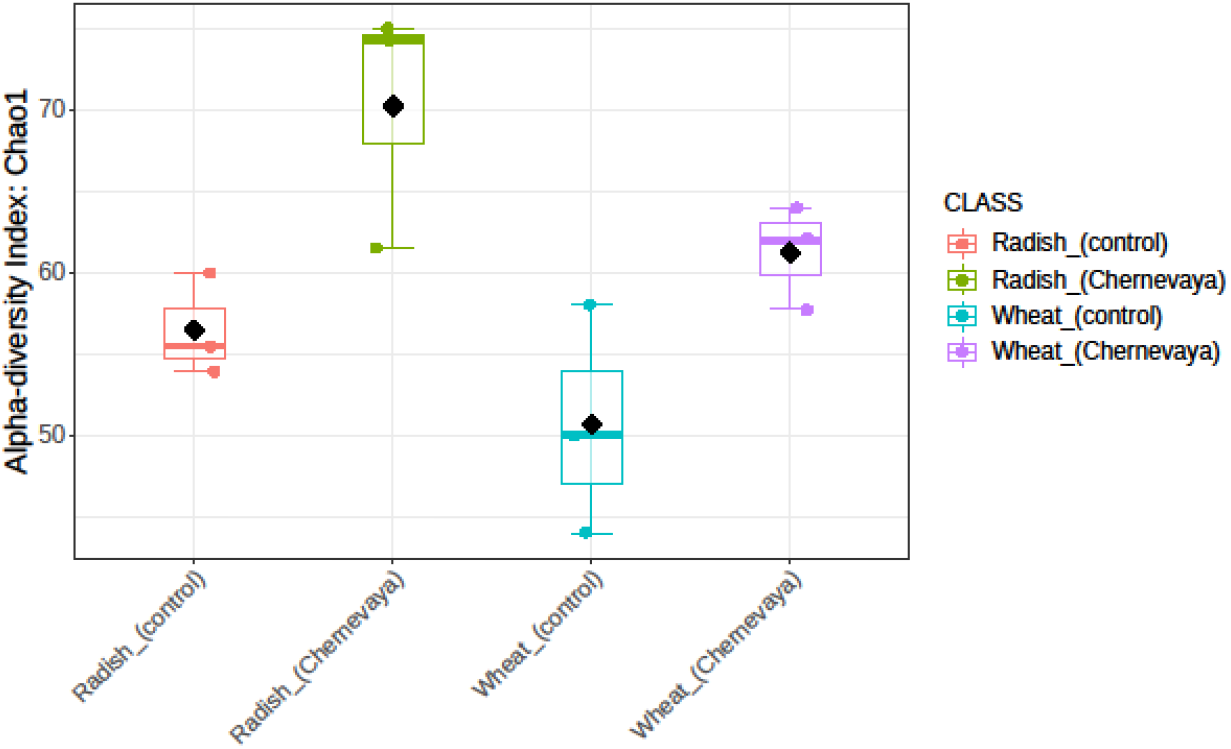
Alpha-diversity measure using Chao1 index in the radish and wheat rhizosphere fungal microbiomes of two soils, aggregated by replications

Furthermore, the beta-diversity analysis was conducted to compare the fungal communities from different sampling sites using PCoA based on the Bray-Curtis index (Fig. 5). Our data show a clear separation of the Chernevaya taiga rhizosphere communities from the control samples based on their fungal community compositions. Additionally, the similarity of species composition between communities of different plants highlights the stability of the fungal communities in the Chernevaya taiga. At the same time, in the control soils, the difference between rhizosphere fungal communities of different plants is much stronger (Fig. 5).

**Fig. 5.**
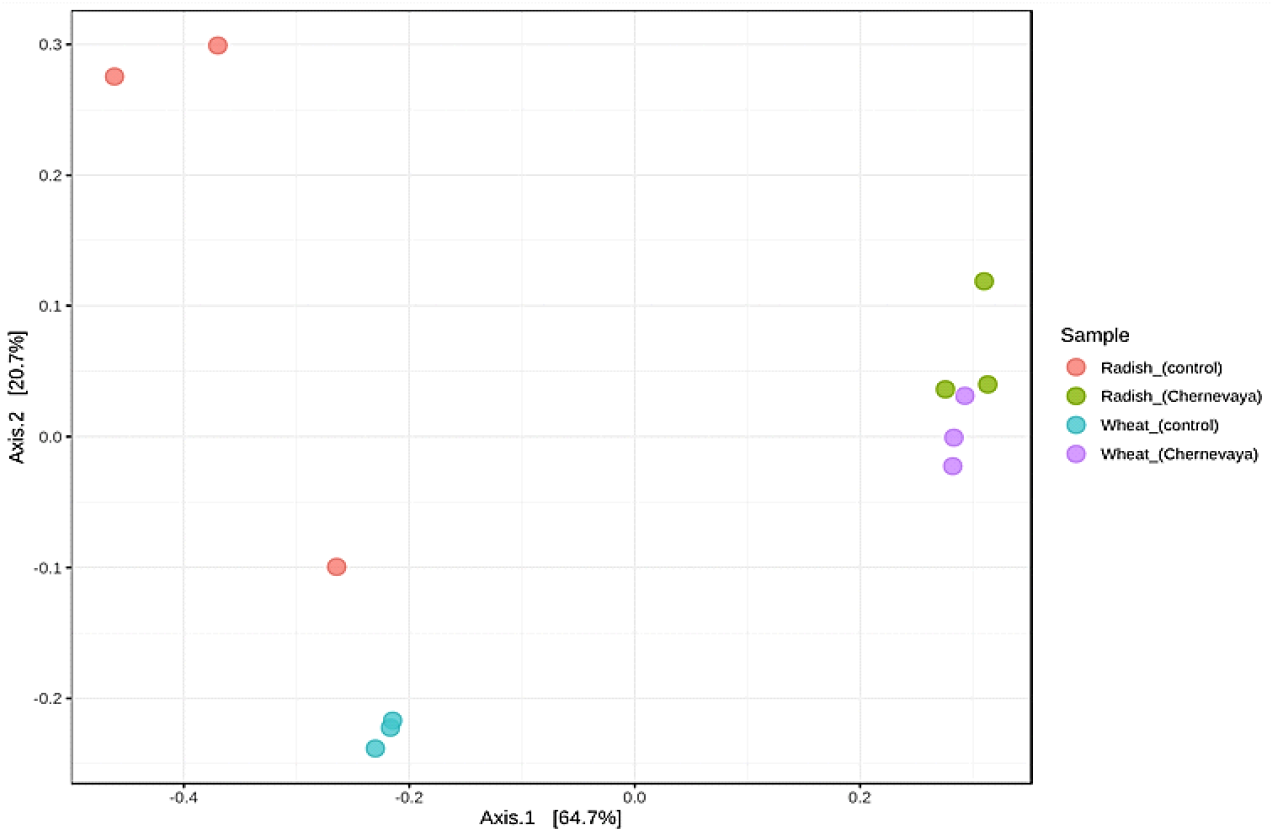
PCoA plot of rhizosphere soil of the radish and wheat plants, grown on the chernevaya and control soils.

### Potential biocontrol and pathogenic fungi in the rhizosphere of crops

We analyzed the distribution of the genera *Trichoderma, Humicola* and *Minimedusa* which are known as biocontrol agents against plant pathogens. Also, we studied the known plant pathogenic fungal taxa (*Fusarium, Oidiodendron*).

It was found that potentially plant-pathogenic fungi were present only in the rhizosphere communities of plants grown in the control soil. In the wheat rhizosphere community, the proportion of *Fusarium* was 4% and that of *Oidiodendro*n was 0.24%. In the rhizosphere of radish, the proportion of *Fusarium* and *Oidiodendron* was much lower, at 0.19% and 0.07%, respectively. At the same time, in the rhizosphere of plants grown on Chernevaya soil T1, representatives of the *Fusarium* genus were absent, while potentially antagonistic fungi were abundant: *Trichoderma harzianum* (4.23–3.91%), *Minimedusa polyspora* (0.04–1.16%), *Humicola fuscoatra* (0.31–1.11%), *H. grisea* (0.34–1.08%), and *Clonostachys ros*ea (0.99–0.11%) (Table 2).

**Table 2.** Abundance of the potential plant biocontrol and pathogenic fungal taxa in the rhizosphere samples.

| Taxa | Raddish - Taiga | Raddish - Control | Wheat - Taiga | Wheat – Control |
| --- | --- | --- | --- | --- |
| Potential biocontrols |  |  |  |  |
| *Trichoderma asperellum* | 0.00% | 2.81% | 0.00% | 3.13% |
| *T.harzianum* | 4.23% | 0.00% | 3.91% | 0.00% |
| *Humicola fuscoatra* | 0.31% | 0.00% | 1.11% | 0.00% |
| *H. grisea* | 0.34% | 0.00% | 1.08% | 0.00% |
| *Minimedusa polyspora* | 0.04% | 0.00% | 1.16% | 0.00% |
| Potential pathogens |  |  |  |  |
| *Fusarium sp.* | 0.00% | 0.19% | 0.00% | 4% |
| *Oidiodendron sp.* | 0.00% | 0.07% | 0.00% | 0.24% |

### Ecological guild analysis

To assess the overall changes in the ecological properties of the studied communities, the FUNGuild approach was used. In general Saprotroph-Symbiotrophic fungi dominate in the rhizosphere of plants grown in Chernevaya taiga soil, while Saprotroph fungi dominated in the rhizosphere community of plants grown in the control soil (Fig. 6).

**Fig. 6.**
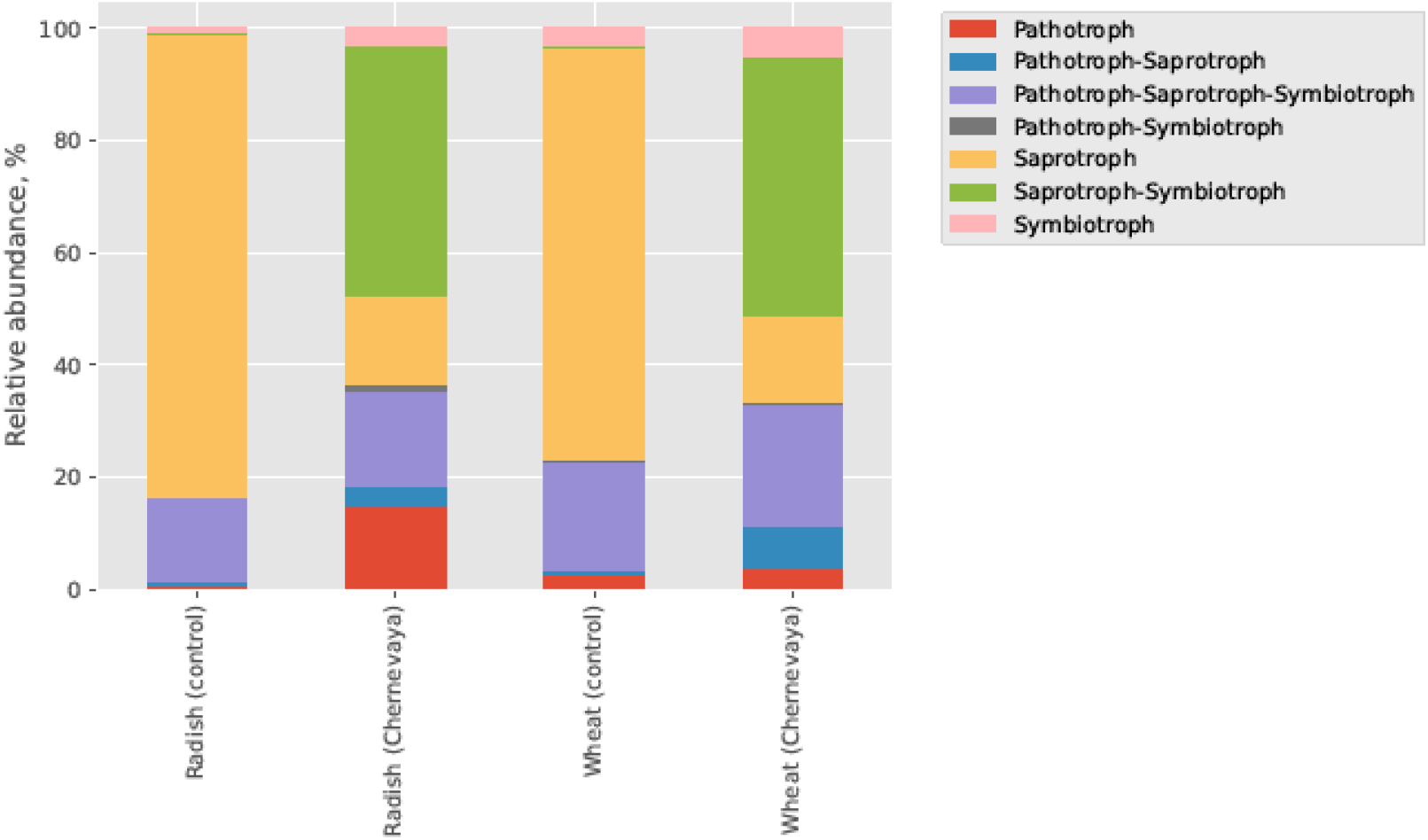
Relative abundance of fungi with various trophic modes in the rhizosphere of wheat and radish growing on the chernevaya and control soils

Regardless of the type of plant combination, a group of fungi with a mixed type of nutrition (Pathotroph-Saprotroph-Symbiotroph) is widely represented in all variants, which is apparently associated with high ecological plasticity. The fraction of Pathotrophs and Symbiotrophs in rhizosphere fungal communit is low. The largest number of Pathotrophs is observed in the rhizosphere of radish grown in the soil of Chernevaya taiga (Fig. 64). At the same time, symbiotrophic fungi are present in the wheat rhizosphere of both soil variants, while in radish, they are present only in the chernevaya soil (T1). It is important to note the proportion of black yeast *Knufia* in this group, which is found among the endophytes of plants growing successfully in extreme conditions (Li et al. 2018).

A less extensive group of Pathotroph-Saprotroph is presented only in the samples from Chernevaya taiga soils. The species *Phialophora finlandia* identified in the rhizosphere of radish grown on T1 soil occurs in nature in ectomycorrhizal and endomycorrhizal associations with conifers. A group of Pathotroph-Symbiotroph is presented in the rhizosphere of wheat grown on the control soil T3, and radish grown on the chernevaya soil T1. This group of fungi is represented mainly by *Oidiodendron chlamydospor*. It is interesting to note that this species forms ericoid mycorrhiza with coniferous plants, and the type materials were isolated from boreal forest soil (Hambleton and Currah 1997).

### Characterization of fungal isolates from rhizosphere of crops growing on the chernevaya soil

In total, 39 strains of fungi were isolated from rhizosphere samples of agricultural crops. To our knowledge, cultivable fungi were assayed in the rhizosphere on crops grown on Chernevaya taiga soil for the first time.

The isolates were divided into morphotypes and identified using morphological and molecular methods. The dominant fungi genera belonged to *Penicillium* (36%, n= 14), *Trichoderma* (23%, n= 9) and *Clonostachys* (21%, n= 8) (Fig. 7).

**Fig. 7.**
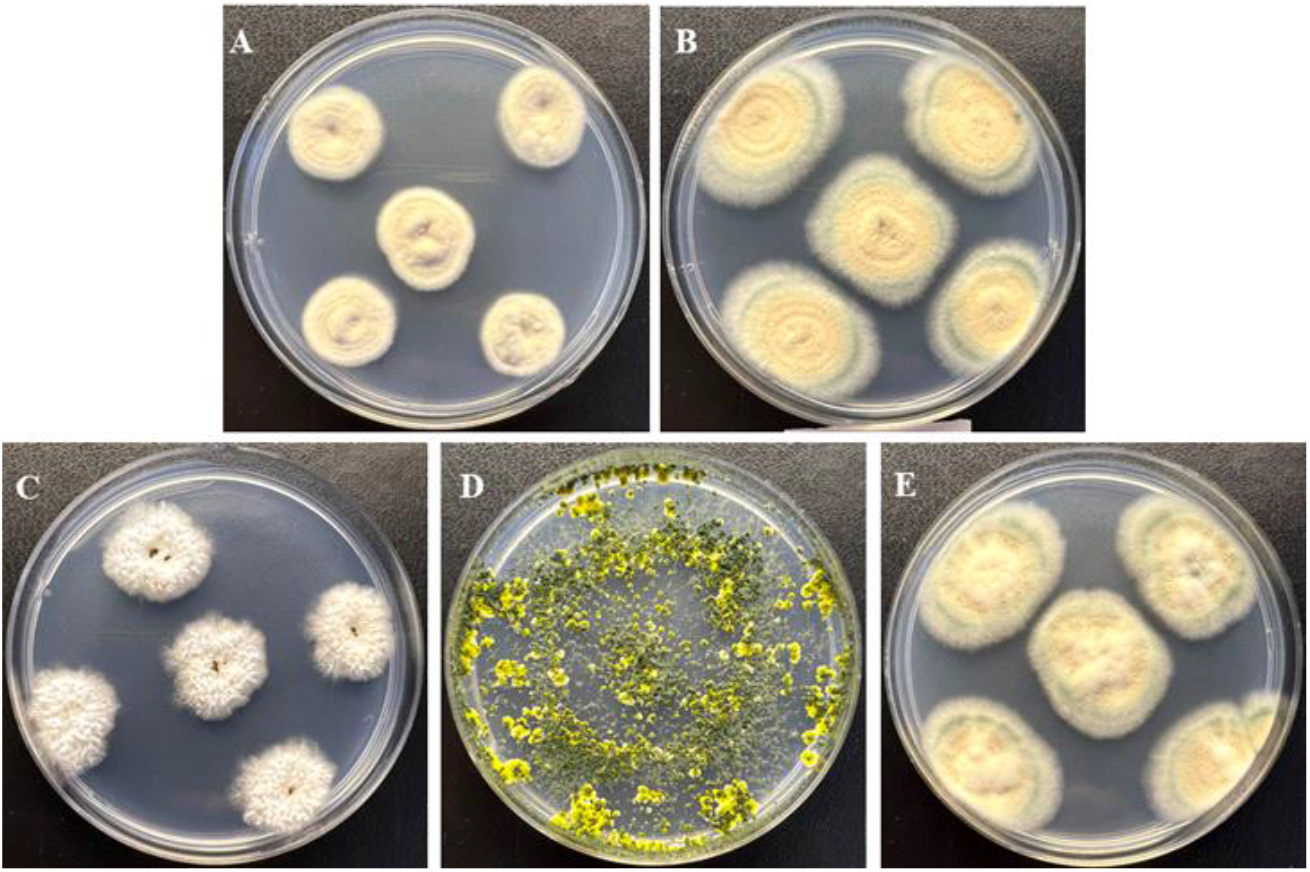
Pure fungal cultures isolated from the rhizosphere soil samples: *Penicillium pancosmium* RR4F (a); *P. camponotu*m RR5F (b); *Clonostachys rosea f. catenulata* SR10F(c); *Trichoderma inhamatum* WH6F (d), *P. camponotum* WH12aF (e).

Isolates from different morphotypes were randomly selected to perform rDNA analysis and phylogenetic identification (Table 3).

**Table 3.** Results of identification of cultivated rhizospheric fungi using ITS region sequence analysis.

| Strain ID (#GenBank) | Nearest relative | Query Cover, % | Identity, % |
| --- | --- | --- | --- |
| RR4F (OQ621732.1) | *Penicillium pancosmium* CBS 276.75 ITS region; from TYPE material (NR_121506.1) |  | 99.46 |
| RR5F (OQ621733.1) | *Penicillium camponotum* CBS 140982 ITS region; from TYPE material (NR_158823.1) | 97 | 98.35 |
| WH12aF (OQ621736.1) | *Penicillium camponotum* CBS 140982 ITS region; from TYPE material (NR_158823.1) |  | 98.54 |
| SR10F (OQ621734.1) | *Clonostachys rosea* f. *catenulata* CBS 154.27 ITS region; from TYPE material (NR_165993.1) |  | 99.81 |
| WH6F (OQ621735.1) | *Trichoderma inhamatum* rRNA genes and ITS1 and ITS2 DNA strain CBS 273.78 (Z68187.1) | 95 | 96.17 |

Fungal isolates were tested for plant-growth stimulation in experiments with wheat seedlings. The results demonstrated that the five isolates significantly promoted wheat seedling growth compared to the control (Table 4). Based on the phenotypic observations, all isolates caused an obvious increase in root length (except WH12aF), shoot height, and dry weight. To the greatest extent, *Trichoderma inhamatum* WH6F has increased the dry mass of the wheat seedlings and shoot height, *Penicillium cam*ponotum WH12aF and *Clonostachys rosea* SR10F stimulated shoot height, and *Penicillium pancosmium* RR4F and *P*.*camponotum* RR5F increased root length. It was shown that *P. pancosmium* RR4F and *Trichoderma inhamatum* WH6F suppressed the growth of *Fusarium oxysporiu*m, a well-known plant pathogen, and RR4F also possesses proteolytic activity (data not presented).

**Table 4.** Influence of the isolated fungi on the wheat seedlings growth.

| Plant growth parameters | RR4F | RR5F | WH12aF | SR10F | WH6F | <sup>1</sup>K | <sup>2</sup>GA<sub>3</sub> |
| --- | --- | --- | --- | --- | --- | --- | --- |
| Root length, mm | 39.64±5.32 | 51.80±2.66 | 31.92±3.51 | 38.41±2.86 | 47.96±3.27 | 34.78±5.72 | 61.74±6.27 |
| Shoot height, mm | **85.60±7.26** | 78.80±4.92 | 81.33±4.87 | **84.18±7.21** | **88.22±6.41** | 54.04±6.36 | 81.35±8.93 |
| Dry mass, mg plant<sup>-1</sup> | **270.5±13.5** | **260.0±13.0** | 230.4±11.5 | 230.1±11.5 | **240.7±12.0** | 180.3±9.1 | 240.2±12.0 |

<sup>1</sup>K- plants without inoculation; <sup>2</sup>GA<sub>3</sub> – treatment with gibberellic acid solution. Values that are greater than those in the GA<sub>3</sub> variant are shown in bold. Means are average ±standard deviations (SD)

The plant growth effect of fungal isolates was compared with the effect of gibberellic acid, a natural microbial derived substance that is used in agriculture as a plant growth regulator to stimulate cell division and elongation of leaves and stems. It was found that the effect of inoculation was comparable and, in some cases, exceeded that for GA3 application regarding the shoot height (Table 4).

The measurements showed that the isolates significantly (p < 0.05) affected the growth and biomass accumulation of wheat seedlings compared to the control (Figure 8). Inoculation with all isolates resulted in an increase in the shoot height to 45.82-63.26% and the dry mass to 27.62-50.03%. The effect on the length of the roots was less pronounced, 10.44-48.93%, and inoculation with WH12aF isolate, exhibited a negative effect.

**Fig. 8.**
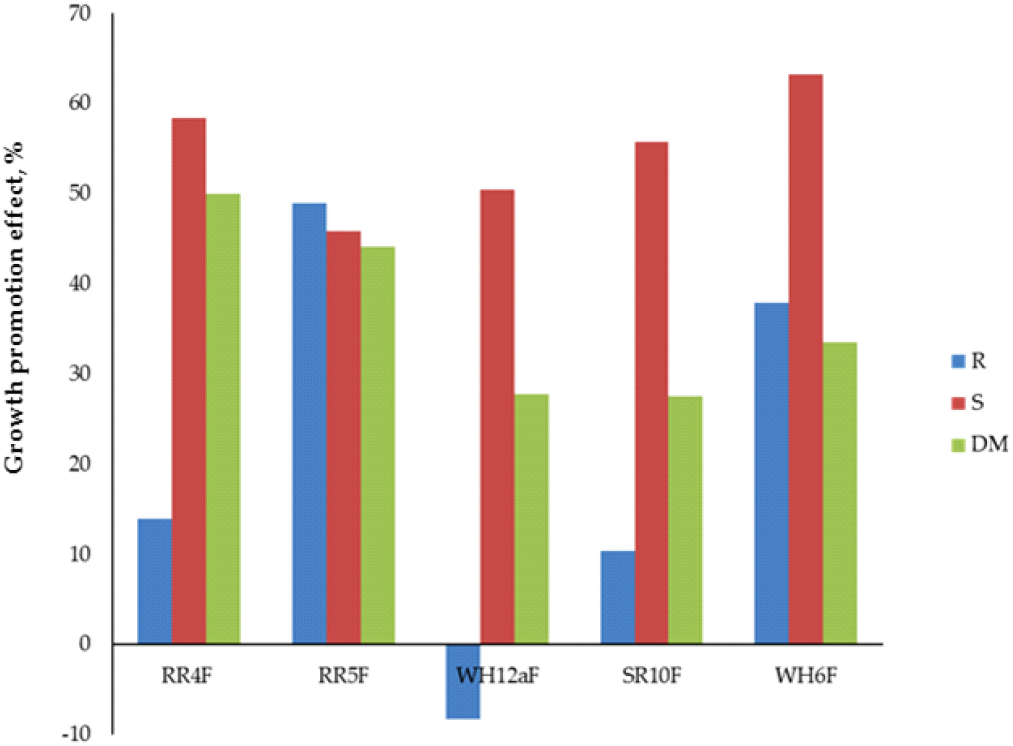
Growth promotion effect of fungal isolates on different growth parameters of the wheat seedlings after 30 days of inoculation. Abbreviations: R-roots length, S-shoot height, DM -dry mass

## Discussion

Chernevaya taiga is characterized by exceptionally high plant productivity. The reasons for this have not yet been definitively determined, but most likely there is a rare combination of some abiotic (soil-forming rocks, climate specificity, soil absorbing complex) and biotic (soil microbial communities) factors, that leads to the phenomenon of gigantism in plants. It was shown that transferring the soil of Chernevaya taiga outside the region of its formation led to a gradual change in its valuable properties. So, the changes in soil bacterial and fungal communities that occur as a result of soil transfer to other conditions are of great scientific and practical interest. From a practical point of view, microbial communities formed in the rhizosphere of agricultural crops grown on the soil of the Chernevaya taiga may be of particular importance.

In the last decade, numerous studies have investigated the composition of root-associated microbial communities. Most of these studies have used next-generation sequencing of microbial marker genes like 16S rRNA for bacteria and the nuclear ribosomal internal transcribed spacer (ITS) region for fungi, while others have used shotgun metagenomics sequencing, where all DNA present in an environmental sample is sequenced. Currently, there are a large number of studies on rhizosphere fungal communities in agricultural crops such as *Arabidopsis, Arabis alpina*, poplar, and sugarcane, and it has been shown that the phyla Ascomycota, Basidiomycota, and less Zygomycota, and Glomeromycota dominate the root mycobiota (de Souza et al. 2016; Bergelson et al. 209; Liu et al. 2023). The high representation of selected fungal phyla in the rhizosphere of different plants suggests that members of these phyla are competitive and adaptable colonizers in various soil types and locations (Boddy and Hiscox 2016; Mueller et al. 2020). Understanding the drivers of the rhizosphere fungal community’s formation and identifying their components is crucial to increasing productivity and reducing pathogen attacks on agricultural crops. Studies aimed at investigating the fungal community of agricultural crops grown in the virgin Chernevaya taiga soil with its extremely high productivity may be a good chance to fill this study gap.

We hypothesized that the composition of fungal rhizospheric communities is closely related to plant productivity and that there is specificity in the composition of communities associated with soil and plant characteristics. The studies have confirmed our assumptions. Significant differences in the composition of rhizospheric fungi have been established for plants grown in chernevaya T1 and control T3 soils. Despite the fact that radish and wheat belong to different families with specific physiological and biochemical characteristics, in the rhizosphere of both plants in T1 soil, the dominant phyla were Ascomycota and Mortierellomycota and a high degree of unidentified species, while in T3 Mucoromycota were the most abundant. These results are in line with the previous metagenome analysis of fungal communities in the chernevaya taiga soils. The most abundant class of fungi in the chernevaya taiga soils were Mortierellomycetes, comprising only the *Mortierella* genus, followed by Agaricomycetes and Globeromycetes (Rayko et al. 2021). The saprotrophic *Mortierella* species are also characteristic of the soils of the middle taiga zone in West Siberia (Filippova and Bulyonkova 2017).

Probably, fungi of this genus can play an important role in soil processes and the stimulation of plant growth due to their characteristics of promoting plant growth and supporting the defense mechanisms in plants. It was demonstrated that most of the species of this genus are able to produce and accumulate in the mycelium polyunsaturated fatty acids (arachidonic, γ-linolenic, eicosapentaenoic, and docosahexaenoic), which are involved in the induction of resistance to phytopathogens in plants of agricultural importance (Zlotek and Wójcik 2014). Another important feature of *Mortierell*a is phosphorus-solubilizing capacity due to the production of several organic acids (Sang et al. 2022). Recent studies revealed other features of *Mortierella* as PGPF, such as the production of phytoregulators (IAA, gibberellic acid, ACC deaminase), siderophores production, strong cellulolytic and chitinolytic activity, protease and urease activity, etc. (see review Ozimek and Hanaka 2021 for details).

Another important discovery of this study is that the yeast *Solicoccozyma* represented a significant part of the fungal communities in the rhizosphere of crops grown in the chernevaya soil. These yeasts are typical soil-borne microorganisms that are commonly isolated from soils worldwide (Yurkov et al. 2012) and also detected in environmental studies (Lynch and Thorn 2006). The predominance of *Solicoccozyma* species in the rhizosphere may be attributed to the unique polysaccharide capsules surrounding those that aid in the assimilation of nutrients from soil and thus compete with bacteria and other fungi (Yurkov 2018).

To determine the contribution of different plant species to the formation of their rhizosphere fungal community in a particular soil, we compared their taxonomic composition, highlighting common and unique taxa. The fungal communities of the rhizosphere soils of wheat and radish grown in T1 soil were close to each other, while they were very different in crops grown in T3 soil (Fig. 5). So, the influence of plant-related factors on rhizosphere fungal communities was much less than to soil type.

It seems logical that representatives of wood-decaying mushroom-forming Agaricomycetes fungi were not found in the rhizosphere of agricultural crops. Representatives of AMF are not only widespread in the soils, but are a general constituent of rhizosphere microflora and are associated with over 70% of the world’s vascular plant species in nearly all terrestrial habitats (Brundrett 2009). It was suggested in our previous work (Rayko et al. 2021) that the AMF play a crucial role in the Chernevaya taiga soil fertility, increasing nutrient and water availability. In the current work, we found a small amount of AMFs only in the wheat rhizosphere, and an absence of AMFs in control soils and the radish rhizosphere. This is an expected result, since plants of the Brassicaceae family do not form symbiosis with AMF (Glenn et al. 1985). We found that in laboratory conditions, the effect of Chernevaya taiga soil on wheat growth was significantly more pronounced than on radish. It is possible that the functions performed by AMF in native conditions have moved to other members of the fungal community. We can assume that in man-made conditions, some ascomycetes may be able to switch to an endophytic lifestyle due to the lack of leaf litter.

The role of rhizosphere fungi in plant disease control is of great interest and importance. Understanding the rules that drive the formation of a plant microbiome and identifying its components is crucial to increasing productivity and reducing pathogen attacks. Plants produce and exude via their roots various metabolites that can affect soil microorganisms, and different rhizodeposits influence the rhizosphere microbiome composition. Under natural conditions, in undisturbed soil, the groups of beneficial and harmful microorganisms remain in a state of dynamic equilibrium, but agricultural use can change that. It is important to mention that the well-known pathogenic fungus *Fusarium* was detected only in the rhizosphere of plants on T3 soil. The absence of *Fusarium* in the fungal community of T1 soil may be explained by the presence of the potential antagonistic fungi *Trichoderma, Humicola, Minimedusa. Trichoderma* species can be parasites and antagonists of many phytopathogenic fungi, thus protecting plants from disease (Vinale et al. 2008). *Humicola* species are considered potential antagonists for the biological control of plant diseases (Ko et al. 2011). It is interesting to note that the composition of biocontrol microorganisms differed in the fungal rhizosphere communities of plants grown on chernevaya and control soil. *Humicol*a and *Mininedusa* were found only in T1 soil, and *Trichoderma* close to different species (*T. asperellum* in T3 soil and *T. harzianum* in T1 soil) were detected. Therefore, antagonistic activity related to phytopathogens depends on specific ecological conditions.

The availability of fungal cultures would be a valuable tool to investigate the functionality of the fungal communities and reconstruct specific rhizosphere fungal communities in vitro. For this purpose, cultivable fungi from the rhizospere of crops grown on T1 soil were considered in this work. Altogether, there are 39 fungal species, which is less than 0.2% of the total fungal diversity obtained by DNA metabarcoding. The isolates were mainly from the Ascomycota division, and different *Penicillium* and *Trichoderma* were the dominant genera in cultivated communities. We have not been able to obtain isolates of *Mortierel*la, which is dominant according to the results of molecular analysis. These findings are in line with data from another study of Chernevaya soil cultivable fungi (Kirtsideli et al. 2022). In general, results based on culture-dependent and molecular techniques are in good agreement. The reasons for differences in the evaluation of fungal community diversity may be because of the difficulty of direct DNA extraction from spores.

One of the important parameters in the development of biorational products for plant protection is the stability of the microbial community. The study of the taxonomic composition of the rhizosphere microbiota of agriculturally significant plants grown in Chernevaya taiga soil under laboratory conditions, together with the previously obtained data (Rayko et al.2021; Kravchenko et al. 2022), opens the way for the creation of sustainable communities in vitro those contribute to the successful development of agricultural crops. Gaining an understanding of the entire structure of the natural community of microorganisms, present in the soil and rhizosphere of the Chernevaya taiga, including bacteria and fungi, can be an important step in the development of nature-like biofertilizers.

To expand our understanding and obtain more complete data on the impact of the fungal communities on plant fertility, it is necessary to expand the range of agricultural crops due to the large influence of host plants on the composition and structure of the rhizosphere microbiota.

## Author contribution

All authors contributed to the study concept and design. Experimentation was performed by IK, ET, AT. The first draft was written by I.K., M.R., S.S. and A.L., and all authors commented on the previous version of the manuscript. All authors read and approved the final manuscript.

## Funding

This research is funded in part by the Saint Petersburg State University (grant ID 94030965). The data analysis was carried out using computational resources provided by the Resource Center “Computer Center of SPbU”. This work was funded by the Russian Science Foundation, grant number 19-16-00049 (field work, 16S rRNA genes amplification, sequencing), by Saint Petersburg State University, grant ID PURE 93023187 (DNA sequencing analyses, phylogenetic analysis) and by the Ministry of Science and Higher Education of the Russian Federation, state task registration number 122040800164-6 (microbiological analyses, pot experiments, isolation, and analysis of fungal isolates).

## Acknowledgments

The study was carried out using the equipment of the resource center “Genomic Technologies, Proteomics and Cell Biology” of ARRIAM and core facilities “Bioengineering” and “UNIQEM collection” of Research Center of biotechnology RAS. The authors are extremely grateful to Prof. Dmitrii Vlasov, Saint Petersburg State University, Department of Botany, for reading and discussing this manuscript and for insightful remarks.

## Data Availability

The sequence data presented in this study are openly available in Figshare at https://doi.org/10.6084/m9.figshare.22639870.v1 (accessed on 14 April 2023). All other data generated or analysed during this study are included in this article.

## Conflict of interests

The authors declare that they have no conflict of interest.

## Notes

### Competing Interest Statement

The authors have declared no competing interest.

